# TCR repertoire analysis reveals effector memory T cells differentiation into Th17 cells in rheumatoid arthritis

**DOI:** 10.1101/616441

**Authors:** Xu Jiang, Shi-yu Wang, Chen Zhou, Jing-hua Wu, Yu-hao Jiao, Li-ya Lin, Xin Lu, Bo Yang, Wei Zhang, Xin-yue Xiao, Yue-ting Li, Xun-yao Wu, Xie Wang, Hua Chen, Li-dan Zhao, Yun-yun Fei, Hua-xia Yang, Wen Zhang, Feng-chun Zhang, Hui Chen, Jian-min Zhang, Bin Li, Huan-ming Yang, Jian Wang, Wei He, Xue-tao Cao, De-pei Liu, Xiao Liu, Xuan Zhang

## Abstract

The pathogenesis of rheumatoid arthritis (RA), a systemic autoimmune disease characterized by autoreactive T-cell accumulation and pro-inflammatory cytokine overproduction, is unclear. Systematically addressing T-cell receptor (TCR) repertoires of different CD4+ T-cell subsets could help understand RA pathogenesis. Here, peripheral CD4+ T cells from treatment-naïve RA patients and healthy controls were sorted into seven subsets including naïve, effector, central memory, effector memory (EMT), Th1, Th17, and regulatory T cells. T-cell receptor β chain repertoires were then analyzed by next-generation sequencing. We identified T-cell clonal expansion in EMT and Th17 cells, with highly similar TCR repertoires between them. Ex vivo experiments demonstrated the preferred differentiation from EMT to Th17 cells in RA. Moreover, TCR diversity in subsets including Th17 was negatively correlated with RA disease activity indices such as C-reactive protein and erythrocyte sedimentation rate. Thus, shared and abnormally expanded EMT and Th17 TCR repertoires might be pivotal for RA pathogenesis.

## Introduction

Rheumatoid arthritis (RA) is a systemic immune disorder that mainly affects joints, causing chronic inflammatory damage with a high rate of disability; however, its etiology and underlying immunological mechanisms are still elusive. Disease modifying anti-rheumatic drugs (DMARDs) such as methotrexate have long been administered in clinical practice and are defined as first-line treatments according to the most recent guidelines^1^. However, these treatments can cause severe adverse effects due to their broad-range activities on the whole organism beyond the immune system.

To overcome the shortcomings of classical DMARDs and to pursue more precise treatment options, accumulating studies have demonstrated that T-cell-mediated immunity plays a pivotal role in the pathogenesis of RA, which could potentially be developed as a drug target^2^. Nevertheless, the efficacy of different T-cell-targeting therapies currently applied in clinical practice remains uncertain^3,4^. Hence, a more thorough understanding of T cell biology in the setting of RA is urgently needed.

Based on basic T-cell biology, it is known that during the process of T-cell activation, CD4+ T cells recognize major histocompatibility complex-presented peptide fragments by expressing rearranged T-cell receptors (TCRs). Some evidence supporting a role for CD4+ T cells in RA pathogenesis is the association between this disease and HLA-DRB1, which encodes the polymorphic HLA class II DRβchain, and especially HLA-DR4 subtypes^5,6^. The autoantigen-driven expansion of effector T-cell clonotypes is crucial for autoimmune inflammatory responses. Therefore, elucidating the TCR repertoire under conditions of autoimmunity could help to uncover the interaction between T-cells and pathogenic antigens in RA.

Previous studies on the TCR repertoire in RA patients were completed via V-D-J gene sequencing and have not yet reached concordant conclusions. Early in 1991, Howell et al. first investigated TCR heterogeneity in RA and claimed that a dominant TCR is present^7^. Later, in 1993, Jenkins et al. discovered accumulation of the Vβ-encoding gene in RA patients^8^. However in 1995 and 1996, together with his own findings, Alam et al. reviewed several articles with controversial conclusions and proposed several possibilities that could cause such discrepancies, such as sample numbers and PCR techniques used for quantitative purposes, among others^9,10^. Thereafter, studies utilizing similar PCR repertoire sequencing technology were performed. Two groups pointed out that TCRs might not accumulate clonally under certain circumstances^11,12^. VanderBorght et al. found that T-cells expand oligoclonally in the synovium, but not in the peripheral blood^11^. Cantaert et al discovered that in anti-citrullinated protein antibody-positive (ACPA+) RA patients, similar preferential TCR clonal selection was observed, compared to that in ACPA− RA patients. Notwithstanding these controversial findings, most studies have demonstrated that the T cells from RA patients respond through clonal expansion in the synovia and/or peripheral blood^11,13,14,15,16,17,18,19,20,21^.

All of these early studies investigated TCR clonality at the pan-T-cell level and inevitably overlooked the differential functions of different subsets, due to limitations in basic techniques and the knowledge of T-cell biology. It is now certain that different T-cell subsets have very distinct roles in initiating and exacerbating the disease. Recent studies showed that some unique T-cell subpopulations, such as Th1 cells and IL-17 producing Th17 cells, are expanded during RA progression. Th1 cells produce interferon (IFN)-γ and tumor necrosis factor (TNF)-α, promoting inflammation in the synovia during RA^22,23^. Further, the Th-17–IL-17–IL-23 axis was found to be crucial to trigger local inflammation and it also contributes to chronicity in murine models^24,25^. Another important regulatory mechanism in RA involves CD4+CD25+ Treg cells, which help to maintain peripheral tolerance and protect against the initiation of autoimmune-mediated pathology^26^. Researchers believe that in this disease, there is an imbalance between Treg and Th17 cells, which leads to unresolved inflammation^27,28^. In other human autoimmune disease studies, Th17 cells were suggested to interact or develop connections with various cell types including Treg, effector memory T cells (EMT), and dendritic cells, among others^29,30,31,32^. However, how these T-cell subsets are derived and how their function is maintained are still unclear, and the aforementioned findings clarify the contention that both pre-existing cytokines and cell–cell interactions lead to disease chronicity.

As described, there is a missing link between T-cell subsets with distinct functions and their contribution to the pan-TCR repertoire. Accordingly, in the present study, we classified CD4+ T cells into naïve (NT), effector (ET), central memory (CMT), and EMT subsets according to antigen experience and cell development processes, in addition to Th1, Th17, and Treg cells, which have been suggested to be involved in the pathogenesis of RA. We utilized TCR repertoire sequencing technologies and bioinformatics strategies to quantitatively evaluate the TCR repertoires of these T-cell subsets from RA patients in an attempt to determine the origins of different subsets and provide clues for novel therapeutic strategies.

## Results

### Sample set

Peripheral blood mononuclear cells (PBMCs) from twelve RA patients and eight age- and gender-matched healthy controls (HCs) were isolated and sorted into seven CD4+ T-cell subsets by flow cytometry, including NT (CCR7+CD45RA+CD45RO−), ET (CCR7−CD45RA+CD45RO), EMT (CCR7−CD45RA−CD45RO+), CMT (CCR7+CD45RA−CD45RO+), Th1 (CD45RA−CD161−CXCR3+CCR6−), Th17 (CD45RA−CD161+CXCR3−CCR6+) and Treg (CD45hi CD127−/lo) (Supplementary Figure S1a and b). RA patients had a significantly increased frequency of Th17 cells and a decreased frequency of Treg as expected. Moreover, RA patients appeared to have fewer CMT cells and a higher but statistically insignificant frequency of EMT cells compared to those in HCs (Supplementary Figure S2). The abnormal expansion of Th17 or EMT cells could possibly be related to RA pathogenesis. Using next-generation sequencing technology, we examined the T-cell receptor beta (TCRB) chain repertoires of different CD4+ T-cell subsets. The RNA yield from cells per subset correlated with the number of cells (Pearson’s R = 0.46; p < 0.00001). From more than 6.5 × 10^6^ CD4+ T-cells, greater than 2.6 × 10^7^ reads and 1.0 × 10^6^ unique CDR3s clonotypes were generated from the data normalized to cell numbers.

### Abnormal gene usage and CDR3 length in RA

The rearrangement of V- and J-gene segments determines the response of CDR3 to HLA complex and antigenic peptides^33^. To evaluate the differences in TRBV and TRBJ repertoires between RA and HC groups, we examined the distribution of TRBV and TRBJ segments of different CD4+ T cell subsets. Among them, NT, EMT, and Th17 subsets demonstrated consistent tendencies regarding the use of V3-1 and J2-7 segments. In RA patients, the use of the TRBJ2-7 gene family was significantly reduced in NT and EMT subsets, and a decreasing trend was observed in the Th17 subset, compared to those in HCs (Fig. 1a). However, the TRBV3-1 gene was associated with higher use in the NT, EMT, and Th17 subsets (Fig. 1b) in RA patients compared to that in HCs. We also detected the decreased use of TRBV6-2 and TRBV6-3 in the NT subset, TRBV15 in the EMT subset, and TRBV7-9 in the Th17 subset (Fig. 1b). Moreover, the use of TRBV3-1, TRBV6-1, TRBV6-2, TRBV6-3, and TRBV6-5 in CMT cells, TRBV24-1, TRBV28, and TRBV7-2 in ET cells, and TRBV12-4 in Tregs was also significantly different in RA patients compared to that in HCs (Supplementary Figure S3). Except for TRBJ2-7, no other J gene expression was significantly different in each subset, when comparing RA to HC groups (Supplementary Figure S4).

**Fig. 1.**
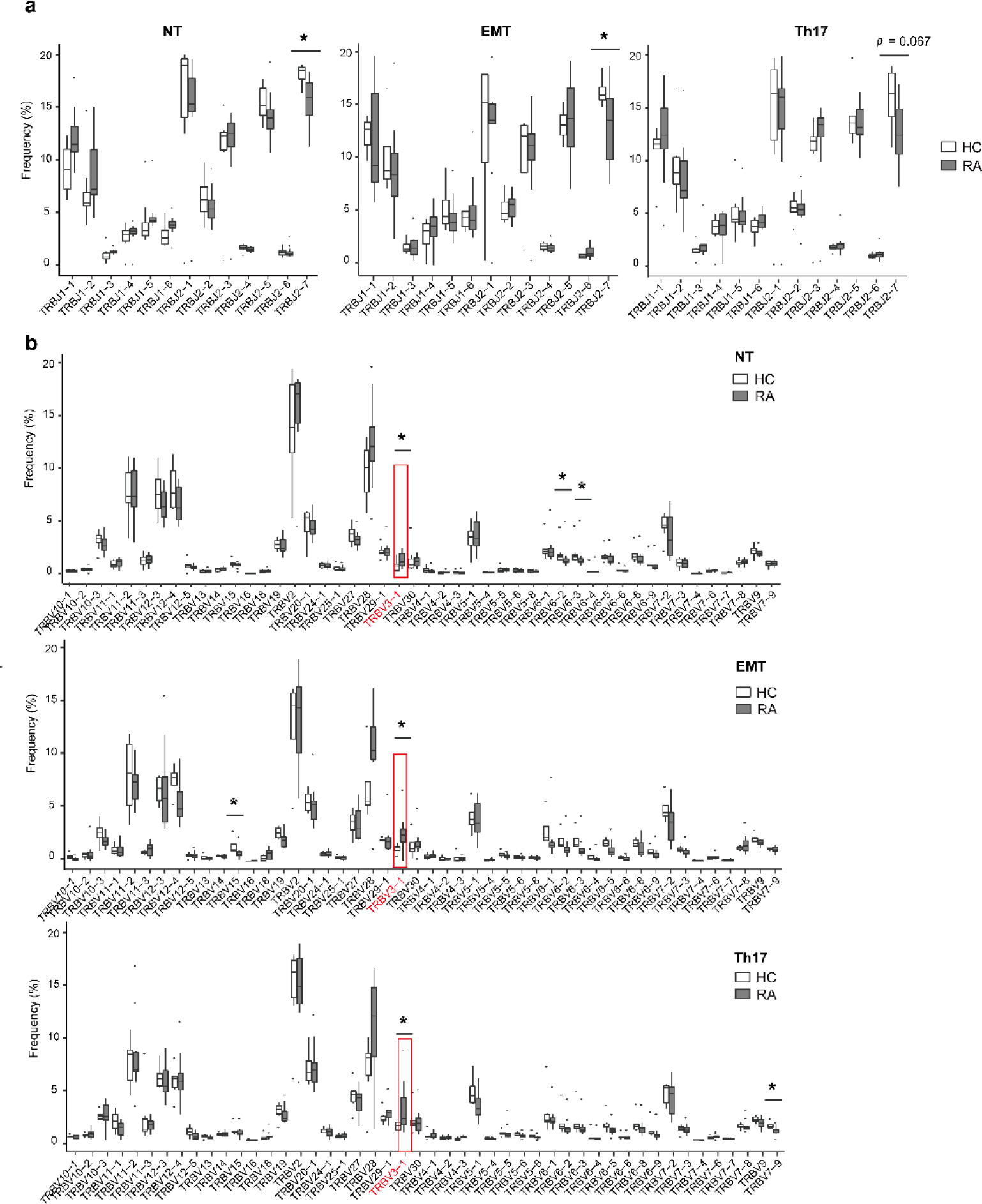
Rheumatoid arthritis (RA) patients show different usages of V/J genes compared to those in healthy controls (HCs). Frequencies of V (a) and J (b) genes used by NT, EMT, and Th17 subsets in RA and HC groups are shown (NT: HC n = 8, RA n = 12; EMT: HC n = 4, RA n = 11; Th17: HC n = 5, RA n = 11; Mann-Whitney U-test; *p < 0.05). NT, naïve T cells; EMT, effector memory T cells.

A previous study on type 1 diabetes found that the length of the TCRB CDR3 was reduced in patients^34^, which resulted in the generation of an enriched repertoire of auto-reactive TCRs and susceptibility to disease pathogenesis. Thus, whether RA, as another typical autoimmune disease, is associated with this phenomenon was of profound interest. CDR3 length changes can be attributed to pre-selection, negative and positive selection in the thymus, antigen selection, or any combination. We therefore examined these processes in detail, by analyzing the length of unproductive (out-of-frame, OOF) TCRB CDR3 in NT cells and productive (in-frame) TCRB CDR3 in different CD4+ T-cell subsets. We found that the OOF CDR3 length in NT cells was differentially distributed (p = 0.018) in RA patients compared to that in HCs (Supplementary Figure S5a). However, the in-frame CDR3 length was similar between RA and HC groups for each CD4+ T-cell subset (Supplementary Figure S5b). As OOF CDR3 would not be affected by thymus selection and NT cells are independent of antigen selection, we concluded that the pre-selected TCR repertoire in RA was biased in CDR3 length, which might be related to RA pathogenesis.

### TCRB repertoires of EMT and Th17 cells are clonally expanded in RA patients

Diversity is an effective measurement to assess the state of the immune system challenged by autoimmune disease, and TCRB diversity in memory T cells was previously reported to be reduced with RA^35,36^. To specify the roles of different cell subsets and to systematically address the change in TCR repertoires in RA, the Shannon-index was used to examine the diversity of each subset, and normalized Diversity 20 (D20) and Diversity 50 (D50) indexes were also calculated to determine clonality. The Th17 subset from RA patients had a lower D20 index compared to that of HCs and the same tendency was observed for the D50 index, suggesting that Th17 cells in RA are more clonal (Fig. 2a and Supplementary Figure S6). The EMT subset from RA had a significantly lower Shannon-index, indicative of less diversity, whereas diversity and clonality in CMT cells were similar between RA and HC groups (Fig. 2a and b). Therefore, Th17 and EMT cells, rather than the other subsets, experienced significant clonal expansion in RA patients. As self-antigens are regarded as key factors in the induction of RA, antigen-specific expansion was expected in Th17 cells from RA patients. In addition, the tendency of decreased EMT diversity in RA patients suggested that these cells could also be influenced by auto-antigens.

**Fig. 2.**
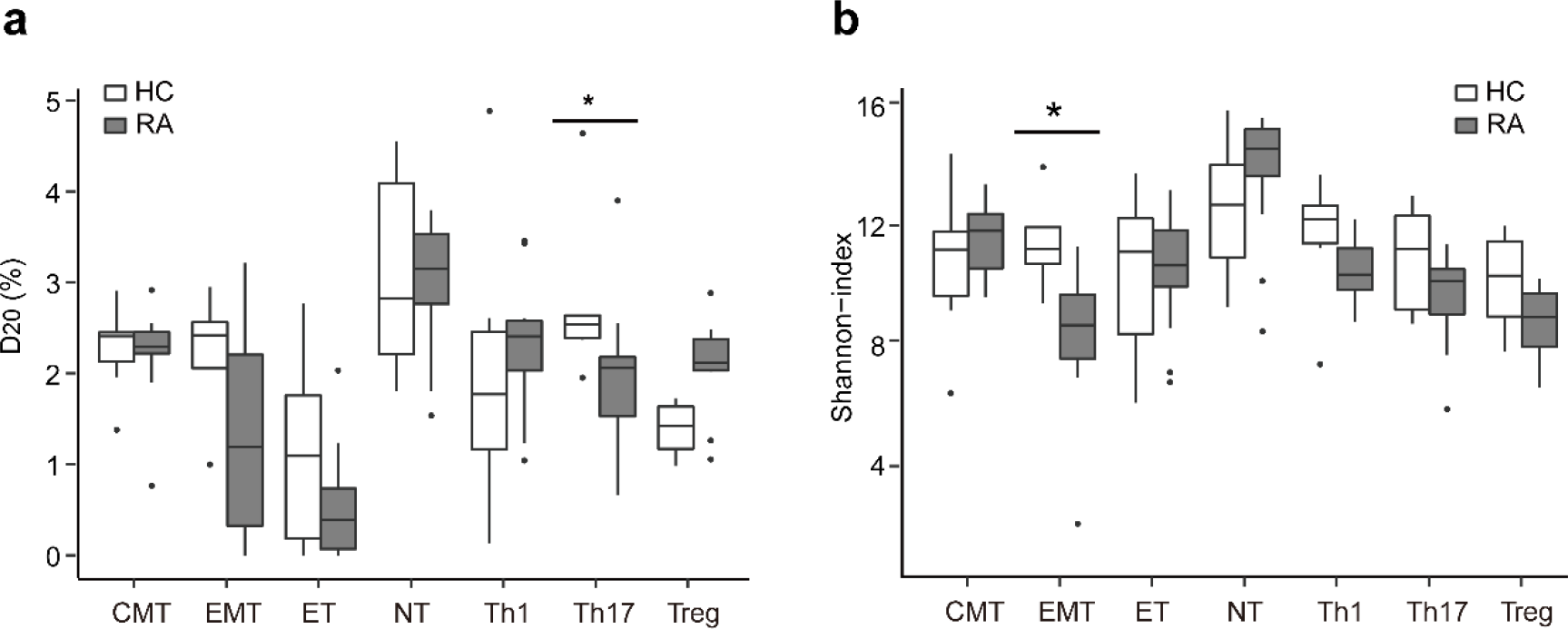
Th17 and effector memory T (EMT) cells from rheumatoid arthritis (RA) patients exhibit less diversity than those from healthy controls (HCs). The D20 index (a) and Shannon-index (b) were calculated; Th17 cells from patients showed less clonality than those from HCs and the Shannon-index of EMT cells showed that the diversity of CDR3.AA from EMT cells of RA patients was lower than that in cells from HCs; For a-b, data of all samples that were sequenced successfully, as shown in supplementary table S3, were analyzed. (Mann-Whitney U-test; *p < 0.05). CMT, central memory T cells; EMT, effector memory T cells; ET, effector T cells; NT, naïve T cells; Treg, regulatory T cells.

### TCRB repertoires of Th17 cells show an increased correlation with EMT cells in RA

TCRs, serving as the unique proxies of T-cells, can be used to track their development and differentiation^37^. As the repertoires of Th17 cells from RA patients exhibited a series of abnormal phenotypes including a higher proportion and increased clonality compared to those in HCs, we examined the correlation between TCRB repertories and Th17 and other subsets using overlapping indices. As shown in Fig. 3a, increasing overlapping indices were observed between Th17 and EMT cells in RA patients (Mann-Whitney U-test, p < 0.05). In contrast, only Th17, and no other subsets, showed significantly increased similarity with EMT cells (Supplementary Figure S7a). This result indicated biased differentiation between EMT and Th17 cells in RA. In addition, the frequencies of overlapping clonotypes were significantly higher than those of private clonotypes in most donors (7/11 for both EMT and Th17 cells), implying the antigen-experienced and clonally expanded tendency of the differentiated T cells (Fig. 3b). As these T cells might participate in the pathogenesis of RA and have clinical importance, we sorted overlapping clonotypes between EMT and Th17 cells in all RA patients based on their frequencies in EMT subsets, filtering the clonotypes present in any HC samples. We identified 167 unique CDR3-amino-acid (CDR3.AA) clonotypes in this manner and listed them in Supplementary Table S1. The characterization of these CDR3s revealed that the vast majority of them (126/167) exhibited increased frequencies in Th17 cells compared to those in EMT cells (Supplementary Figure S7b), and their length distribution was significantly increased compared to their 74 counterparts in HCs and the private clonotypes (Fig. 3c). Motif analysis did not reveal any significant differences between RA and HC groups (Supplementary Figure S7c).

**Fig. 3.**
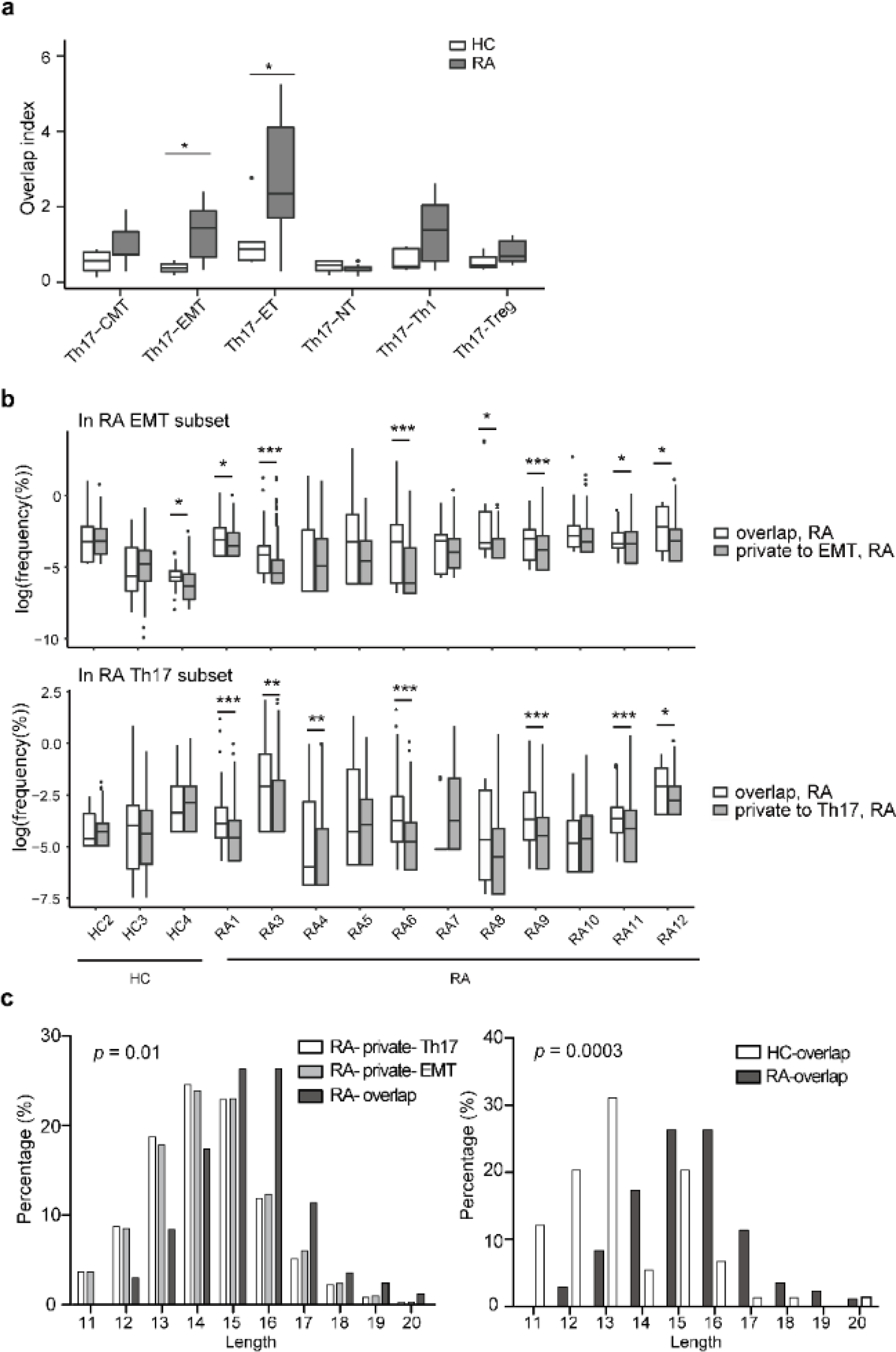
T cell receptor beta (TCRB) CDR3s are highly shared across Th17 and effector memory T (EMT) cells from rheumatoid arthritis (RA) patients. (a) The overlapping index between Th17 and other subsets was calculated and CDR3s of Th17 and EMT cells from RA patients exhibited more similarity compared to those from healthy controls (HCs). (b) In RA, the frequencies of overlaps between EMT and Th17 clonotypes were higher than those of private clonotypes in EMT (up) and Th17 (bottom) subsets. (c) Clonotypes belonging to overlaps between EMT and Th17 (EMT-Th17) cells were longer than private clonotypes (left) and HC EMT-Th17 overlapping clonotypes; For a, the overlap index was calculated between subsets separated from the same donor, as shown in the supplementary table S3. (χ^2^-test with simulated p-value (based on 2000 replicates) and Mann-Whitney U-test; *p < 0.05; **p < 0.01; *** p < 0.001). CMT, central memory T cells; EMT, effector memory T cells; ET, effector T cells; NT, naïve T cells; Treg, regulatory T cells.

### EMT cells in RA patients are inclined to differentiate into Th17 cells *ex vivo*

Th17 cells were found to be crucial for the development of RA; however, the sources and maintenance of specific antigen-expanded Th17 cells are still unclear. Mounting studies support the plasticity of the Th17 lineage; meanwhile, researchers believe that committed Th17 cells can survive the contraction phase to form EMT cells^38,39^. In addition to being differentiated from naïve T cells, Th17 cells can also proliferate and undergo self-renewal or can be induced by memory CD4+ T cells *in vitro*^40,41^. As an increase in the overlapping indices between EMT and Th17 cells was identified in PBMCs from RA patients in this study, the differentiation from EMT to Th17 cells could be another pathway to sustain expanded Th17 subsets in RA. To test this hypothesis, *ex vivo* experiments were conducted to determine if EMT or CMT cells could yield Th17 cells. EMT and CMT cells were sorted from PBMCs of RA and HC groups and then stimulated with anti-CD3/CD28 and cultured in the presence of anti-IL4, anti-INF-γ, and Th17-inducing cytokines (IL-1β, IL-6, IL-23, TGF-β) for 6 days. *In vitro*-induced Th17 cells were detected by the intracellular staining of IL-17A (Fig. 4a). As shown in Fig. 4b, EMT cells from RA patients were more inclined to differentiate into Th17 cells than those from HCs (p < 0.05), whereas this phenomenon was not observed in CMT cells from RA patients.

**Fig. 4.**
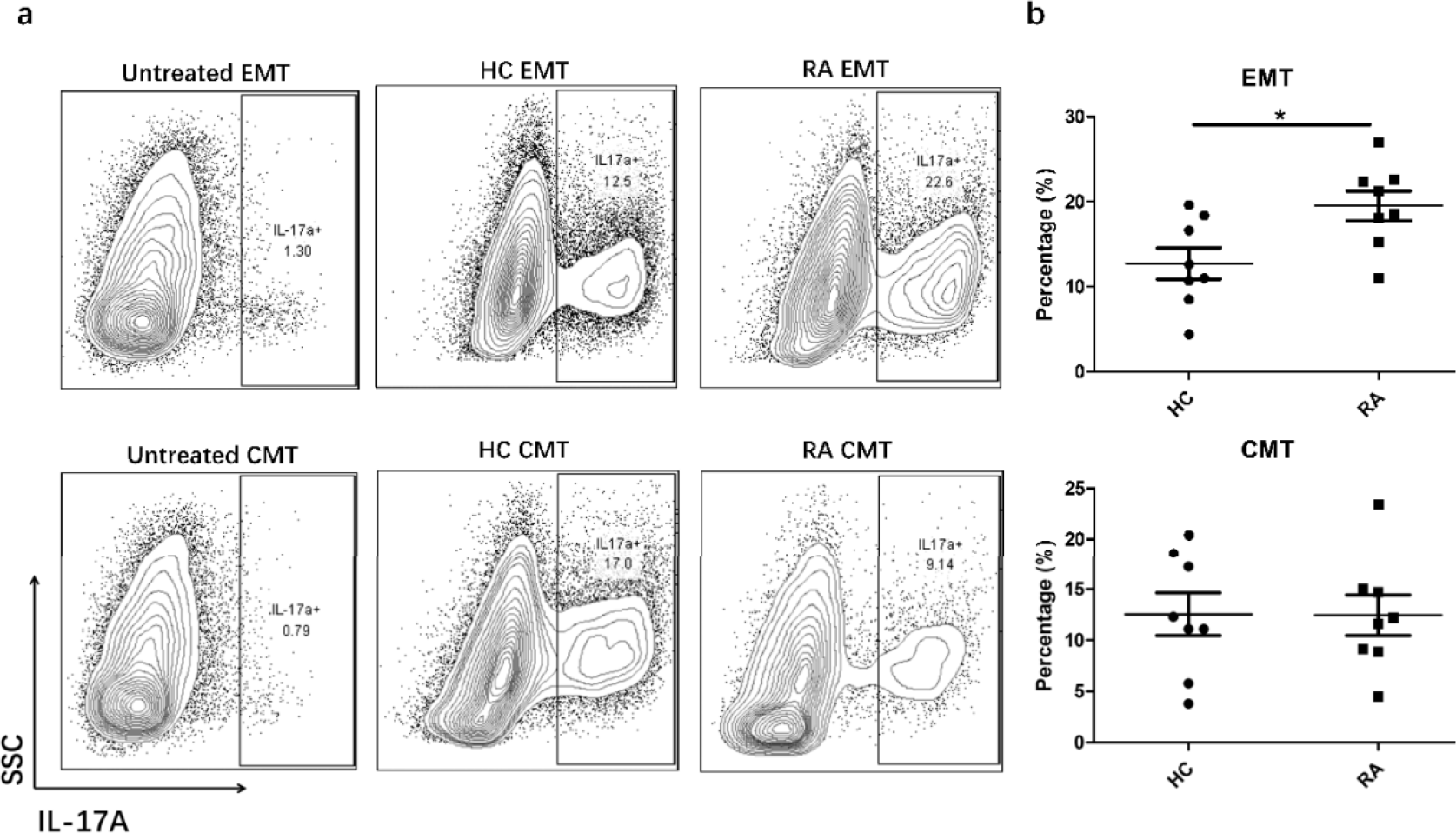
Effector memory T (EMT) cells from rheumatoid arthritis (RA) patients induce more Th17 cells *in vitro*. EMT and central memory T (CMT) cells isolated from peripheral blood mononuclear cells (PBMCs) of RA and healthy control (HC) groups were cultured with 3 μg/ml anti-CD3/CD28, 5 μg/ml anti-IL4, 5 μg/ml anti-INF-γ, Th17-inducing cytokines IL-1β, IL-6, IL-23 (all at 10 ng/ml), and TGF-β (5 ng/ml) for 6 days. Control cells were cultured without these cytokines. (a) Cytokines from treated and untreated cells from RA patients and HCs were collected and the expression of IL-17A was detected by intracellular staining and flow cytometry. (b) Frequencies of Th17 cells induced by EMT and CMT cells from RA patients and HCs *in vitro* (n = 8; Student’s t-test; *p < 0.05).

### TCRB CDR3 clonalities of certain cell subsets, including Th17 cells, correlate with clinical indicators of RA

Clonality analysis of Th17 cells in RA patients showed the biased expansion of some clonotypes. To determine if the accumulation of these clonotypes was related to pathogenesis and/or disease activity, we analyzed the correlation between the D20 index and clinical indicators of RA and found that the D20 index of Th17 cells negatively correlated with C-reactive protein (CRP) (Cor −0.72, p = 0.024; Fig. 5a), erythrocyte sedimentation rate (ESR) (Cor −0.85, p = 0.002; Fig. 5b), and 28-joint disease activity score (DAS28) (Cor −0.52, p = 0.1; Fig. 5c). Furthermore, an inverse correlation between the D20 index of Treg and Th1 cells and serum IgG levels was found in RA patients (Cor −0.89, p = 0.001; Fig. 5d and Cor −0.79, p = 0.006; Fig. 5e, respectively).

**Fig. 5.**
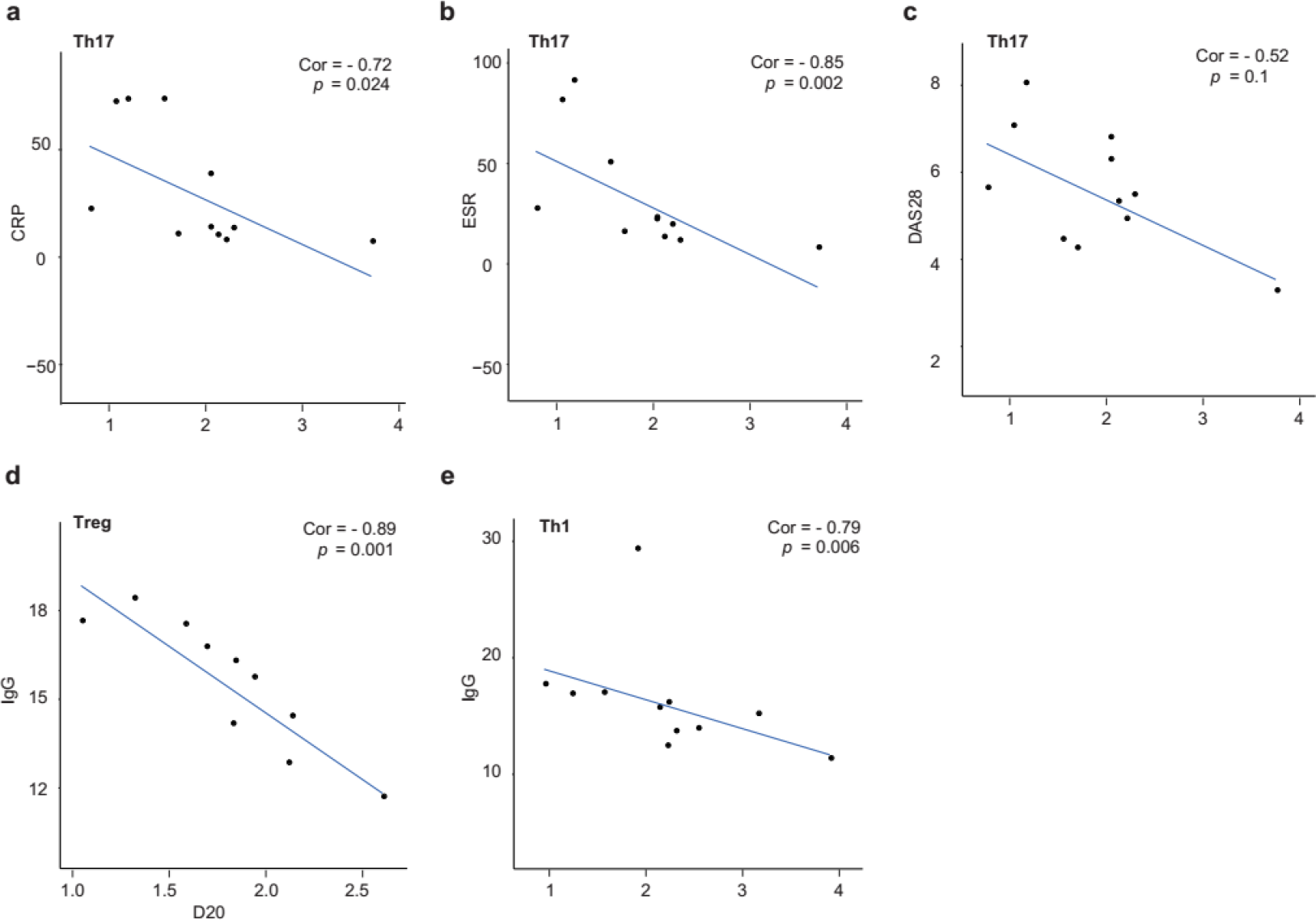
The D20 indexes of the T cell receptor beta (TCRB) of each subset from rheumatoid arthritis (RA) patients negatively correlates with clinical indexes. The D20 index of TCRB of Th17 cells was found to negatively correlate with C-reactive protein (CRP) (a), erythrocyte sedimentation rate (ESR) (b), and the 28-joint disease activity score (DAS28) (c), whereas the D20 index of Treg (d) and Th1 (e) cells negatively correlated with IgG levels (Th17 n = 11; Th1 n = 11; Treg n = 10; Spearman’s correlation).

We then annotated the functions of CDR3 clonotypes from each sample based on the knowledge dataset (https://db.cngb.org/pird/tbadb/), which records disease-related sequences and information on these sequences. As shown in Supplementary Figure S8, T-cells in CMT subsets from RA patients included more pathogen-related clonotypes, suggesting that pathogens could trigger RA^42^. Due to the limited record of auto-antigen-related TCR sequences, the remaining clonotypes unique to RA patients could not be characterized based on a determined immunological explanation.

## Discussion

RA is a complex autoimmune disease with multiple etiological factors, among which, aberrant T-cell activation plays a key role in the initiation and perpetuation of the disease. In the present study, we focused on seven important CD4+ T-cell subsets, namely NT, ET, EMT, CMT, Th1, Th17, and Treg, considering both antigen experience during cell development processes and T-cell immune responses and regulation.

The biased use of the Vα-gene and Vβ-gene has been reported in T cells from synovial fluid; however, the significantly different use of any V-gene was not identified in the peripheral blood^21,43,44^. In this study, we found that the use of 11 V-genes and 1 J-gene by the TCRB was different in certain subsets between RA and HC groups. Notably, among them, we found that use of the V3-1 and J2-7 segments showed a similar trend for NT, EMT, and Th17 subsets. The abnormal use of these two genes has been observed in numerous autoimmune and infectious diseases. The TRBJ2-7 gene was found to be abnormally expressed in patients with systemic lupus erythematosus^45^, type 1 diabetes^46^, and chronic hepatitis B^47^. Alexandra et al. reported that the TRBV3-1 gene was highly expressed in Th1 and Th17 cells of primary Sjögren’s syndrome patients^48^. Further, the abnormal expression of TRBV3-1 and TRBJ2-7 might be associated with the pathogenesis of immune-related diseases. V/J gene use in healthy individuals was suggested to be mostly influenced by HLA^49^, but the antigen-driven effect should not be ignored^50,51^. Commonalities in the usage bias of TRBV3-1/TRBJ2-7 in NT, EMT, and Th17 cells suggested a stronger correlation between these subsets. Further work is needed to explore this relationship. Our study also indicated that the OOF CDR3 length in NT cells was reduced in RA patients, suggesting that early events during T cell development and the re-arrangement of the TCR in RA might be abnormal. We speculate that the auto-reactive TCR repertoire was probably generated during biased early V(D)J gene recombination and that this pre-selected repertoire might contribute to RA susceptibility.

Studies have reported that TCR repertoire diversity is decreased in RA and other autoimmune diseases due to the specific autoantigen-derived amplification of T cell clonality. In this study, we highlighted the decreased diversity and increased clonality of TRB CDR3 in Th17 and EMT cells based on Shannon-index and D20 index analysis, indicating a high frequency of clonal amplification these subsets. Furthermore, we found correlations between different subsets, especially Th17 and EMT cells, in RA and HC groups, which could reflect a common source of these amplified clonotypes. Both the frequencies of common clonotypes and the proportions of unique clonotypes between Th17 and EMT subsets were significantly increased in RA compared to those in HCs. Taken together, in terms of both V/J gene expression and antigen-driven expansion, Th17 and EMT subsets demonstrated abnormal phenotypes and stronger correlations in RA patients.

Th17 cells are involved in numerous autoimmune diseases^52^ and can be activated by IL-6, which is secreted by antigen presenting cells and T cells^53^. The depletion of interferon (IFN)-γ was shown to contribute to Th17-related inflammation and the administration of anti-IL-17A monoclonal antibodies was found to alleviate arthritis in mice^54^. However, in humans, how Th17 cells are regulated and how they contribute to chronic inflammation with RA have not been elucidated^55^. Studies have detected increased levels of cytokines like TGF-β, IL-6, and IL-21 in both the periphery and affected joints^56^. As Th17 cells are required for the generation of activation signals from pro-inflammatory cytokines like TGF-β, IL-6, IL-21, and IL-23, among others^57^, the mechanism underlying the continuous maintenance of Th17 cells and their prolonged activation remains elusive. Collectively, the amplification of Th17 in both the periphery and at the inflammatory site including synovial fluid and the synovium of RA patients has been demonstrated in previous studies. Further TCR sequencing analysis indicated that the amplification of Th17 cells occurs in an oligoclonal manner. However, the source of Th17 subsets or the mechanism underlying their clonal expansion in RA remained unclear. Our study attempted to answer this question. Specifically, increased clonal overlap between Th17 and EMT cells was observed in RA patients compared to that in HCs. Moreover, the overlapping clonotypes were further found to be expanded clonotypes. Consequently, our findings suggested that there is a cohort of expanded oligoclonal Th17 cells that share similar TCR clonotypes with EMT cells. Further, these cells might also belong to the exact EMT lineage that exhibits a shared TCR repertoire, instead of other T-cell lineages such as CMT cells, which was further supported by *ex vivo* experiments. The current results led us to a hypothesis regarding a possible pathogenic mechanism associated with RA. Specifically, we suggest that auto-reactive EMT cells with abnormal CDR3 lengths are expanded upon stimulation in RA patients, and that these are differentiated into Th17 cells to facilitate further downstream immune reactions upon re-stimulation by specific antigens.

The correlation between the TCR clonality of Th17 cells and inflammation and disease activity was consistent with the important role of these cells in RA; accordingly, antibodies targeting IL-17 have been proven to reduce CRP levels and DAS28^58^. The negative correlation between Treg cells and IgG suggested that the decrease in IgG levels stemmed from the suppression of specific clonotypes of Treg cells that proliferate in response to RA. This was consistent with a previous study that identified an inverse correlation between circulating CD4+CXCR5+CD25highCD127low follicular Treg (Tfr) cells in RA patients and serum IgG levels and demonstrated *in vitro* that Tfr cells suppress B-cell IgG secretion in RA patients with stable disease remission^24^(24).

In summary, through analyzing the TCR repertoire of different CD4+T cell subsets in RA patients, we detected the abnormal expression of TRBV3-1 and TRBJ2-7 segments. This loss of diversity in RA patients was mainly found to exist in Th17 and EMT subsets. More importantly, we identified a correlation between Th17 and EMT subsets and the differentiation from EMT cells to Th17 cells in RA patients, which might play a key role in RA pathogenesis.

## Methods

### Patients and controls

This study was approved by the Institutional Review Board of Peking Union Medical College (PUMC) Hospital. Written informed consent was obtained from each RA patient and HCs. RA was diagnosed at the PUMC Hospital according to the 2010 American College of Rheumatology and EULAR classification criteria. All enrolled RA patients were positive for rheumatoid factor (RF) and/or ACPA and were treatment naïve at their first visit; specifically, they had not yet been treated with any of the following agents: DMARDs, glucocorticoid, TNF inhibitors, or any other biological agents or Chinese traditional medicines. Individuals with a history of severe chronic infection, any current infection, severe underlying physical disorders such as acute coronary syndrome, chronic renal failure, or any type of tumor were not included in this study. Pregnant or lactating women were excluded as well. Laboratory data regarding serum levels of CRP, ESR, RF, ACPA, anti-keratin antibody (AKA), anti-perinuclear factor (APF), and anti-mutated citrullinated vimentin (anti-MCV) were recorded. Disease activity was measured using the DAS28 calculated by CRP (DAS28-CRP). The clinical features of all patients are presented in Supplementary Table S2.

### Sample collection and fluorescence-activated cell sorting (FACS) analysis

Peripheral blood was collected, and PBMCs were isolated by Ficoll-Hypaque density-gradient centrifugation. PBMCs were washed twice with FACS staining buffer (PBS, 5 FBS, 0.1% sodium azide) and stained with antibodies for 40 min on ice. Subsets of CD4+ T cells were sorted into different tubes using a BD FACS Aria II flow cytometer. Fluorochrome-conjugated monoclonal antibodies were used to recognize human CD4, CD197 (CCR7), CD196 (CCR6), CD25, CD45RA, CD45RO, CD161, CD183 (CXCR3), CD127, IL-17A (BioLegend). For the detection of intracellular IL-17A expression, cells were first stimulated with phorbol myristate acetate and ionomycin for 6 h with GolgiStop in complete RPMI-1640 media in an incubator at 37 °C with 5% CO_2_. Stimulated cells were then fixed and permeabilized with a fixation/permeabilization kit (eBioscience) before intracellular staining.

### TCRB repertoire sequencing library construction and sequencing

RNA was isolated from sorted cells using Trizol (Invitrogen) as indicated by the manufacturer’s guidelines. Entire samples of isolated RNA were used to synthesize cDNA, and then the CDR3 regions of the TCRB were enriched by multiplex PCR and sequenced as previously described^59^. In brief, gDNA was removed from RNA samples with DNase I digestion (NEB) for 10 min at 37 °C. Then cDNA was synthesized from purified RNA using SuperScript I reverse transcriptase (Invitrogen) with a primer specific to the TCRB constant region (5′-CACGTGGTCGGGGWAGAAGC-3′). cDNA was used as the template for multiplex PCR to enrich the target regions of CDR3s with a pool of 30 TRBV primers and 13 TRBJ reverse primers. After PCR, the cDNA with a band size of 110–180 bp was extracted from the 2% agarose gel and purified. Paired-end 150-bp sequencing was then performed using the Hiseq 4000 system (Illumina) following the instructions of the manufacturer.

### Alignment and data analysis

Raw sequencing data were analyzed using IMonitor^60^, which included four main steps for alignment and error correction. First, basic quality control was performed to filter out the low-quality reads and then the clean paired-end reads were merged. Second, two-step BLAST alignment was performed on these merged data according to the references of V, D, J germline genes, and alleles from IMGT (http://www.imgt.org/) to select the best V/D/J alignment for each merged read. Third, CDR3s with low abundance (less than 3 reads/10^6^) were filtered. For the optimized analysis of the diversity of similarity, which was influenced by the depth of sequencing, we sampled five reads per cell for normalization. At last, multiple analyses were performed with IMonitor and further bioinformatics analyses were conducted using R (v3.4.1) (Supplementary Figure S9). The diversity and clonality of samples were evaluated based on the Shannon-index, D20 index, and D50 index. With normalized reads, the Shannon-index was calculated as follows:

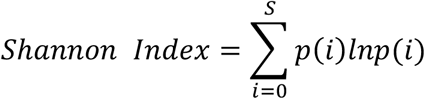

where S is the sum of unique CDR3.AA sequences, which is regarded as a clonotype and *p*(*i*) indicates the frequency of the *i*th clonotype. D20 and D50 represent the percentage of dominant unique clonotypes accounting for 20% and 50% of the total CDR3.AA sequences in a given sample, respectively; the D20 and D50 values were normalized to the total number of clonotypes^61^.

### Overlap indices

Jaccard indices were used to quantify shared clonotypes between two samples from different subsets from one donor as follows:

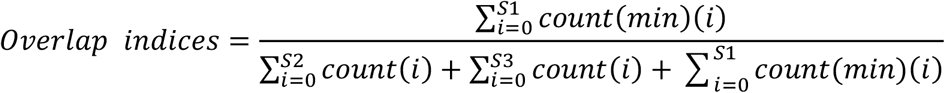

S1 is the sum of number of overlaps; the *i*th overlapping clonotype might have different numbers of sequences for each sample; *count*(min)(i) refers to the smaller value; S2 and S3 represent the sum of the number of clonotypes within each sample, respectively.

### Motif analysis of CDR3s amino acids

Motif analysis was calculated using MEME suite 5.0.3 (http://meme-suite.org/)^62^.

### Statistics

In-house R scripts and GraphPad Prism were employed for this study. To compare the differences between two groups, the two-tailed student’s t-test or two-tailed Mann-Whitney U-test was used. The Spearman method was used to calculate correlations and the χ^2^-test was used to determine the length distribution statistic of overlapping clonotypes. P values < 0.05 were considered significant.

## Supporting information

Supplementary_all

## Data availability

The sequence data that support the findings of this study have been deposited in the China Nucleotide Sequence Archive (CNSA) with the primary accession code CNP0000357.

## Acknowledgments

This study was supported by grants from the National Natural Science Foundation of China (81788101, 81630044, 81601432, 81550023, 81325019, 81771763, 81273312, 91542000, 81801633, 81701623), Chinese Academy of Medical Science Innovation Fund for Medical Sciences (CIFMS2016-12M-1-003, 2017-12M-1-008, 2017-I2M-3-011, 2016-12M-1-008), National Key Research and Development Program: Precise Medical Research (2016YFC0903900), Grant from Medical Epigenetics Research Center, Chinese Academy of Medical Sciences (2017PT31035), Shenzhen Municipal Government of China (JCYJ20170817145404433, JCYJ20170817145428361), Construction Project of National Traditional Chinese Medicine Clinical Research Base of SATCM, and Clinical Cooperative Project of Chinese and Western Medicine for Major and Knotty Diseasesd of SATCM.

## Authors contributions

X.Z. and X.L. designed and supervised the study; C.Z., X.J., X.L., B.Y., X.W., H.C., L.Z., Y.F., H.Y., W.Z., and F.Z. collected samples and performed clinical analysis; X.J., C.Z., Y.J., and X.X. designed and performed experiments; S.W., J.W., W.Z., B.L., H.Y., and J.W. analyzed and interpreted the data; L.L. and X.W designed and performed experiments for sequencing; X.J., S.W., X.Z. X.L., C.Z., J.W., Y.J., and Y.L. drafted the manuscript; H.C., J.Z., W.H., X.C., and D.L. revised the manuscript.

## Reference

1 Singh, J. A. et al. 2015 American College of Rheumatology Guideline for the Treatment of Rheumatoid Arthritis. Arthritis Care Res (Hoboken) 68, 1–25, doi:10.1002/acr.22783 (2016).

2 B., M. I. & G., S. The pathogenesis of rheumatoid arthritis. New England Journal of Medicine 365, 2205–22019 (2011).

3 Bykerk, V. Unmet needs in rheumatoid arthritis. J Rheumatol Suppl 82, 42–46, doi:10.3899/jrheum.090131 (2009).

4 Isaacs, J. D. Therapeutic T-cell manipulation in rheumatoid arthritis: past, present and future. Rheumatology (Oxford) 47, 1461–1468, doi:10.1093/rheumatology/ken163 (2008).

5 Gregersen, P. K., Silver, J. & Winchester, R. J. The shared epitope hypothesis. An approach to understanding the molecular genetics of susceptibility to rheumatoid arthritis. Arthritis & Rheumatism: Official Journal of the American College of Rheumatology 30, 1205–1213 (1987).

6 Stastny, P. Association of the B-cell alloantigen DRw4 with rheumatoid arthritis. New England Journal of Medicine 298, 869–871 (1978).

7 Howell, M. D. et al. Limited T-cell receptor beta-chain heterogeneity among interleukin 2 receptor-positive synovial T cells suggests a role for superantigen in rheumatoid arthritis. Proceedings of the National Academy of Sciences 88, 10921–10925 (1991).

8 Jenkins, R. N., Nikaein, A., Zimmermann, A., Meek, K. & Lipsky, P. E. T cell receptor V beta gene bias in rheumatoid arthritis. J. Clin. Invest. 92, 2688–2701 (1993).

9 Alam, A. et al. Persistence of dominant T cell clones in synovial tissues during rheumatoid arthritis. J Immunol 156, 3480–3485 (1996).

10 Alam, A. et al. T-cell receptor variable region of the β-chain gene use in peripheral blood and multiple synovial membranes during rheumatoid arthritis. Human Immunology 42, 331–339 (1995).

11 VanderBorght, A., Geusens, P., Vandervyver, C., Raus, J. & Stinissen, P. Skewed T-cell recepor variable gene usage in the synovium of early and chronic rheumatoid arthritis patients and persistence of clonally expanded T cells in a chronic patient. Rheumatology 39, 1189–1201 (2000).

12 Cantaert, T. et al. Alterations of the synovial T cell repertoire in anti-citrullinated protein antibody-positive rheumatoid arthritis. Arthritis Rheum 60, 1944–1956, doi:10.1002/art.24635 (2009).

13 Ikeda, Y. et al. High frequencies of identical T cell clonotypes in synovial tissues of rheumatoid arthritis patients suggest the occurrence of common antigen-driven immune responses. Arthritis & Rheumatism: Official Journal of the American College of Rheumatology 39, 446–453 (1996).

14 Schmidt, D., Martens, P. B., Weyand, C. M. & Goronzy, J. J. The repertoire of CD4+ CD28-T cells in rheumatoi arthritis. Molecular Medicine 2, 608 (1996).

15 Goronzy, J. J., Zettl, A. & Weyand, C. M. T cell receptor repertoire in rheumatoid arthritis. International Reviews of Immunology 17, 339–363, doi:10.3109/08830189809054410 (1998).

16 Wagner, U. et al. Clonally expanded CD4+CD28null T cells in rheumatoid arthritis use distinct combinations of T cell receptor BV and BJ elements. Eur. J. Immunol. 33, 79–84 (2003).

17 Klarenbeek, P. L. et al. Inflamed target tissue provides a specific niche for highly expanded T-cell clones in early human autoimmune disease. Ann Rheum Dis 71, 1088–1093, doi:10.1136/annrheumdis-2011-200612 (2012).

18 Spreafico, R. et al. A circulating reservoir of pathogenic-like CD4+ T cells shares a genetic and phenotypic signature with the inflamed synovial micro-environment. Ann Rheum Dis 75, 459–465, doi:10.1136/annrheumdis-2014-206226 (2016).

19 Rao, D. A. et al. Pathologically expanded peripheral T helper cell subset drives B cells in rheumatoid arthritis. Nature 542, 110–114, doi:10.1038/nature20810 (2017).

20 Rossetti, M. et al. TCR repertoire sequencing identifies synovial Treg cell clonotypes in the bloodstream during active inflammation in human arthritis. Ann Rheum Dis 76, 435–441, doi:10.1136/annrheumdis-2015-208992 (2017).

21 Musters, A. et al. In Rheumatoid Arthritis, Synovitis at Different Inflammatory Sites Is Dominated by Shared but Patient-Specific T Cell Clones. J Immunol 201, 417–422, doi:10.4049/jimmunol.1800421 (2018).

22 Kanik, K. et al. Distinct patterns of cytokine secretion characterize new onset synovitis versus chronic rheumatoid arthritis. The Journal of rheumatology 25, 16–22 (1998).

23 Rönnelid, J. et al. Production of T-cell cytokines at the single-cell level in patients with inflammatory arthritides: enhanced activity in synovial fluid compared to blood. British journal of rheumatology 37, 7–14 (1998).

24 Lubberts, E. The IL-23-IL-17 axis in inflammatory arthritis. Nat Rev Rheumatol 11, 415–429, doi:10.1038/nrrheum.2015.53 (2015).

25 Miossec, P. & Kolls, J. K. Targeting IL-17 and TH17 cells in chronic inflammation. Nature Reviews Drug Discovery 11, 763–776, doi:10.1038/nrd3794 (2012).

26 Lohr, J., Knoechel, B., Nagabhushanam, V. & Abbas, A. K. T-cell tolerance and autoimmunity to systemic and tissue-restricted self-antigens. Immunological reviews 204, 116–127 (2005).

27 Niu, Q., Cai, B., Huang, Z.-c., Shi, Y.-y. & Wang, L.-l. Disturbed Th17/Treg balance in patients with rheumatoid arthritis. Rheumatology International 32, 2731–2736, doi:10.1007/s00296-011-1984-x (2011).

28 Samson, M. et al. Brief report: inhibition of interleukin-6 function corrects Th17/Treg cell imbalance in patients with rheumatoid arthritis. Arthritis Rheum 64, 2499–2503, doi:10.1002/art.34477 (2012).

29 Acosta-Rodriguez, E. V. et al. Surface phenotype and antigenic specificity of human interleukin 17-producing T helper memory cells. Nat Immunol 8, 639–646, doi:10.1038/ni1467 (2007).

30 Liu, H. & Rohowsky-Kochan, C. Regulation of IL-17 in Human CCR6+ Effector Memory T Cells. The Journal of Immunology 180, 7948–7957, doi:10.4049/jimmunol.180.12.7948 (2008).

31 Takeshita, M. et al. Polarization diversity of human CD4+ stem cell memory T cells. Clin Immunol 159, 107–117, doi:10.1016/j.clim.2015.04.010 (2015).

32 van Beelen, A. J. et al. Stimulation of the intracellular bacterial sensor NOD2 programs dendritic cells to promote interleukin-17 production in human memory T cells. Immunity 27, 660–669, doi:10.1016/j.immuni.2007.08.013 (2007).

33 Hughes, M. M. et al. T cell receptor CDR3 loop length repertoire is determined primarily by features of the V (D) J recombination reaction. European journal of immunology 33, 1568–1575 (2003).

34 Gomez-Tourino, I., Kamra, Y., Baptista, R., Lorenc, A. & Peakman, M. T cell receptor β-chains display abnormal shortening and repertoire sharing in type 1 diabetes. Nature communications 8, 1792 (2017).

35 Wagner, U. G., Koetz, K., Weyand, C. M. & Goronzy, J. J. Perturbation of the T cell repertoire in rheumatoid arthritis. Proceedings of the National Academy of Sciences 95, 14447–14452 (1998).

36 Oftedal, B. E. et al. T cell receptor assessment in autoimmune disease requires access to the most adjacent immunologically active organ. J Autoimmun 81, 24–33, doi:10.1016/j.jaut.2017.03.002 (2017).

37 Pacholczyk, R., Ignatowicz, H., Kraj, P. & Ignatowicz, L. Origin and T cell receptor diversity of Foxp3+CD4+CD25+ T cells. Immunity 25, 249–259, doi:10.1016/j.immuni.2006.05.016 (2006).

38 Kryczek, I. et al. Human TH17 cells are long-lived effector memory cells. Science translational medicine 3, 104ra100–104ra100 (2011).

39 Pepper, M. & Jenkins, M. K. Origins of CD4+ effector and central memory T cells. Nature immunology 12, 467 (2011).

40 Colin, E. et al. 1, 25-dihydroxyvitamin D3 modulates Th17 polarization and interleukin-22 expression by memory T cells from patients with early rheumatoid arthritis. Arthritis & Rheumatism: Official Journal of the American College of Rheumatology 62, 132–142 (2010).

41 Baricza, E. et al. Distinct In Vitro T-helper 17 Differentiation capacity of Peripheral naive T cells in rheumatoid and Psoriatic arthritis. Frontiers in immunology 9, 606 (2018).

42 Doran, M. F., Crowson, C. S., Pond, G. R., O’Fallon, W. M. & Gabriel, S. E. Frequency of infection in patients with rheumatoid arthritis compared with controls: a population-based study. Arthritis & Rheumatism 46, 2287–2293 (2002).

43 Bröker, B. M. et al. Biased, T cell receptor V gene usage in rheumatoid arthritis. Oligoclonal expansion of T cells expressing Vα2 genes in synovial fluid but not in peripheral blood. Arthritis & Rheumatism: Official Journal of the American College of Rheumatology 36, 1234–1243 (1993).

44 Jenkins, R. N., Nikaein, A., Zimmermann, A., Meek, K. & Lipsky, P. E. T cell receptor V beta gene bias in rheumatoid arthritis. The Journal of clinical investigation 92, 2688–2701 (1993).

45 Sui, W. et al. Composition and variation analysis of the TCR β-chain CDR3 repertoire in systemic lupus erythematosus using high-throughput sequencing. Molecular immunology 67, 455–464 (2015).

46 Tong, Y. et al. T cell repertoire diversity is decreased in type 1 diabetes patients. Genomics, proteomics & bioinformatics 14, 338–348 (2016).

47 Qu, Y. et al. High-throughput analysis of the T cell receptor beta chain repertoire in PBMCs from chronic hepatitis B patients with HBeAg seroconversion. Canadian Journal of Infectious Diseases and Medical Microbiology 2016 (2016).

48 Voigt, A. et al. Unique glandular ex-vivo Th1 and Th17 receptor motifs in Sjögren’s syndrome patients using single-cell analysis. Clinical Immunology 192, 58–67 (2018).

49 Sharon, E. et al. Genetic variation in MHC proteins is associated with T cell receptor expression biases. Nature genetics 48, 995 (2016).

50 Heather, J. M. et al. Dynamic perturbations of the T-cell receptor repertoire in chronic HIV infection and following antiretroviral therapy. Frontiers in immunology 6, 644 (2016).

51 Sun, J. et al. Composition and Variation Analysis of the T Cell Receptor β-Chain Complementarity Determining Region 3 Repertoire in Neonatal Sepsis. Scandinavian journal of immunology 86, 418–423 (2017).

52 Komatsu, N. & Takayanagi, H. Arthritogenic T cells in autoimmune arthritis. The international journal of biochemistry & cell biology 58, 92–96 (2015).

53 Hirota, K. et al. T cell self-reactivity forms a cytokine milieu for spontaneous development of IL-17+ Th cells that cause autoimmune arthritis. J Exp Med 204, 41–47, doi:10.1084/jem.20062259 (2007).

54 Hickman-Brecks, C. L., Racz, J. L., Meyer, D. M., LaBranche, T. P. & Allen, P. M. Th17 cells can provide B cell help in autoantibody induced arthritis. J Autoimmun 36, 65–75, doi:10.1016/j.jaut.2010.10.007 (2011).

55 Firestein, G. S. & McInnes, I. B. Immunopathogenesis of Rheumatoid Arthritis. Immunity 46, 183–196, doi:10.1016/j.immuni.2017.02.006 (2017).

56 McInnes, I. B. & Schett, G. Cytokines in the pathogenesis of rheumatoid arthritis. Nat Rev Immunol 7, 429–442, doi:10.1038/nri2094 (2007).

57 Zambrano-Zaragoza, J. F., Romo-Martinez, E. J., Duran-Avelar Mde, J., Garcia-Magallanes, N. & Vibanco-Perez, N. Th17 cells in autoimmune and infectious diseases. Int J Inflam 2014, 651503, doi:10.1155/2014/651503 (2014).

58 Liu, C. et al. Increased Circulating Follicular Treg Cells Are Associated With Lower Levels of Autoantibodies in Patients With Rheumatoid Arthritis in Stable Remission. Arthritis & Rheumatology 70, 711–721 (2018).

59 Liu, X. et al. Systematic comparative evaluation of methods for investigating the TCRβ repertoire. PloS one 11, e0152464 (2016).

60 Zhang, W. et al. IMonitor: a robust pipeline for TCR and BCR repertoire analysis. Genetics 201, 459–472 (2015).

61 Tipton, C. M. et al. Diversity, cellular origin and autoreactivity of antibody-secreting cell population expansions in acute systemic lupus erythematosus. Nat Immunol 16, 755–765, doi:10.1038/ni.3175 (2015).

62 Bailey, T. L. & Elkan, C. Fitting a mixture model by expectation maximization to discover motifs in biopolymers. Proc Int Conf Intell Syst Mol Biol 2, 28–36 (1994).

