## Supplementary_all for "TCR repertoire analysis reveals effector memory T cells differentiation into Th17 cells in rheumatoid arthritis"

Table S1 Clinical features of RA patients

| | RA patients |
|---|---|
| Gender (male/female) | 1/11 |
| Age (years) | 44.5±3.8 |
| Disease duration (months) | 49.4±15.8 |
| Morning stiffness (minutes) | 95.8±47.1 |
| Tender joint count (n) | 14.5±2.5 |
| Swollen joint count (n) | 7.6±2.3 |
| DAS28-CRP | 5.5±0.4 |
| ESR (mm/H) | 34.8±9.0 |
| CRP (mg/L) | 29.4±9.0 |
| IgG (g/L) | 15.7±1.5 |
| IgA (g/L) | 2.2±0.2 |
| IgM (g/L) | 1.5±0.3 |
| RF (IU/ml) | 98.7±36.4 |
| RF+ (%)* | 66.7 |
| ACPA+ (%)* | 75 |
| AKA/APF+ (%) | 25 |
| Anti-MCV+ (%)* | 50 |

*RF+ is defined as >20U/ml. ACPA+ is defined as >25IU/ml. Anti-MCV+ is defined as >20IU/ml. The value is shown as mean ± SD.

Table S2. Samples derived from each donor.

| | NT | CMT | EMT | ET | Th1 | Th17 | Treg |
|---|---|---|---|---|---|---|---|
| RA1 | 1 | 1 | NA | 1 | NA | NA | NA |
| RA2 | 1 | 1 | 1 | 1 | 1 | 1 | 1 |
| RA3 | 1 | 1 | 1 | 1 | 1 | 1 | 1 |
| RA4 | 1 | 1 | 1 | 1 | 1 | 1 | 1 |
| RA5 | 1 | 1 | 1 | 1 | 1 | 1 | NA |
| RA6 | 1 | 1 | 1 | 1 | 1 | 1 | 1 |
| RA7 | 1 | 1 | 1 | 1 | 1 | 1 | 1 |
| RA8 | 1 | 1 | 1 | 1 | 1 | 1 | 1 |
| RA9 | 1 | 1 | 1 | 1 | 1 | 1 | 1 |
| RA10 | 1 | 1 | 1 | 1 | 1 | 1 | 1 |
| RA11 | 1 | 1 | 1 | 1 | 1 | 1 | 1 |
| RA12 | 1 | 1 | 1 | 1 | 1 | 1 | 1 |
| HC1 | 1 | 1 | NA | 1 | NA | NA | NA |
| HC2 | 1 | 1 | 1 | 1 | 1 | 1 | 1 |
| HC3 | 1 | 1 | 1 | 1 | 1 | 1 | 1 |
| HC4 | 1 | NA | 1 | 1 | 1 | 1 | 1 |
| HC5 | 1 | 1 | NA | 1 | NA | NA | NA |
| HC6 | 1 | 1 | NA | 1 | 1 | 1 | NA |
| HC7 | 1 | 1 | NA | 1 | 1 | 1 | NA |
| HC8 | 1 | 1 | 1 | 1 | 1 | NA | 1 |

NA: FACS, sequencing failing.

1: Samples which sequenced successfully and of which data was used in analyses.

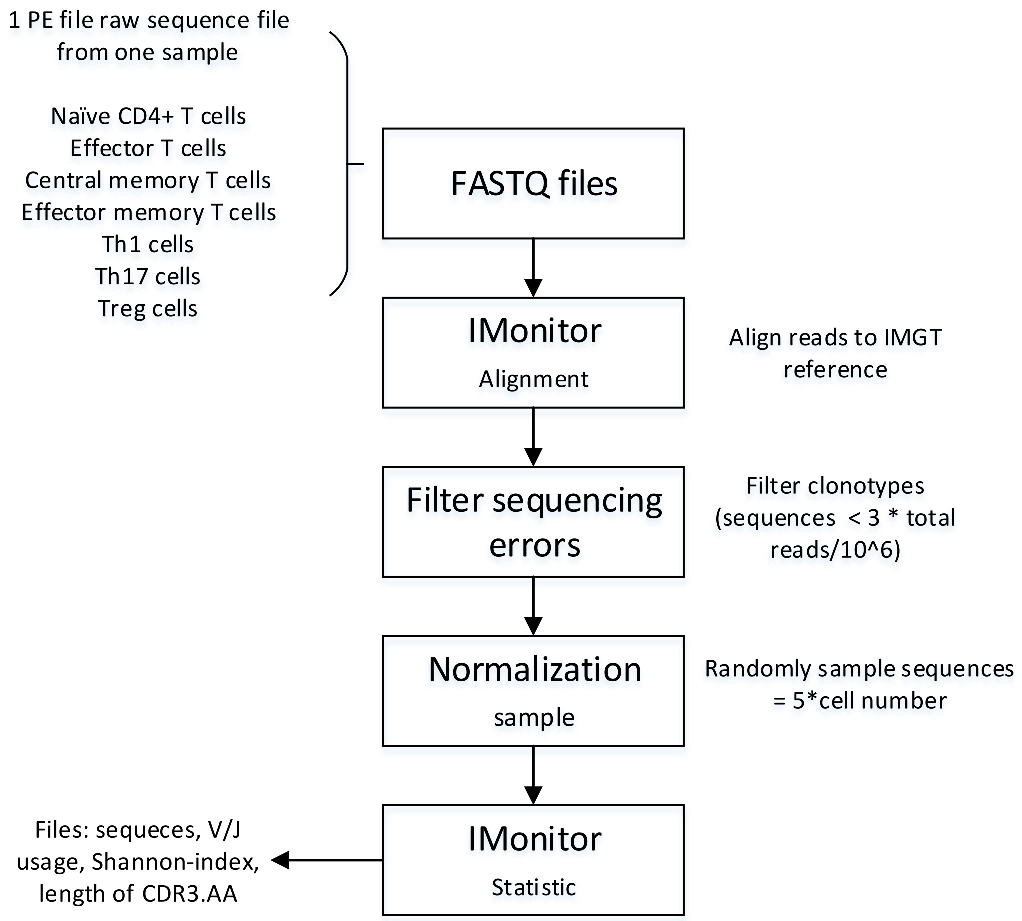


Figure S1 Diagram of data analysis method.

The pipeline for alignment, filter and normalization.

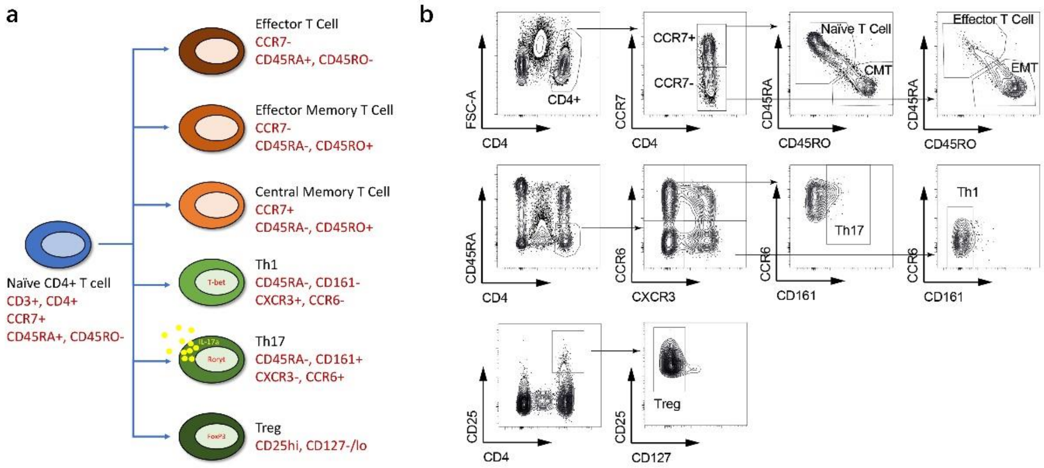


Figure S2 Strategy of the CD4+ T cell subsets sorting by flow cytometry.

(a)  Cell surface markers used by FACS. (b) Staining pattern of each subset.

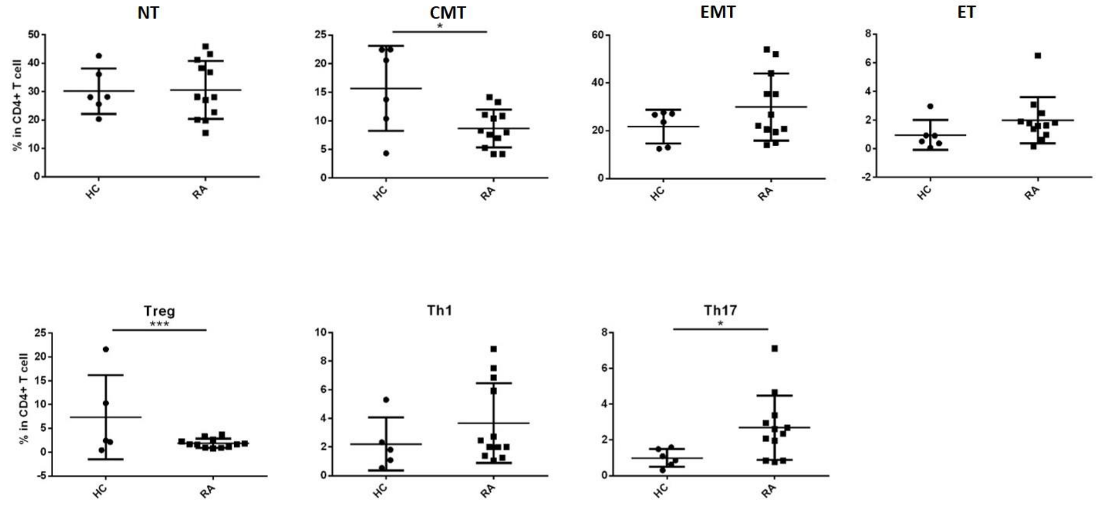


Figure S3 Frequency of each subset in CD4+ T cells in RA patients and HC.

Each subset of RA and HC was sorted by flow cytometry and the frequency of each subset in CD4+ T cell was analyzed. (Unpaired t-test. *p<0.05; ***p<0.001.)

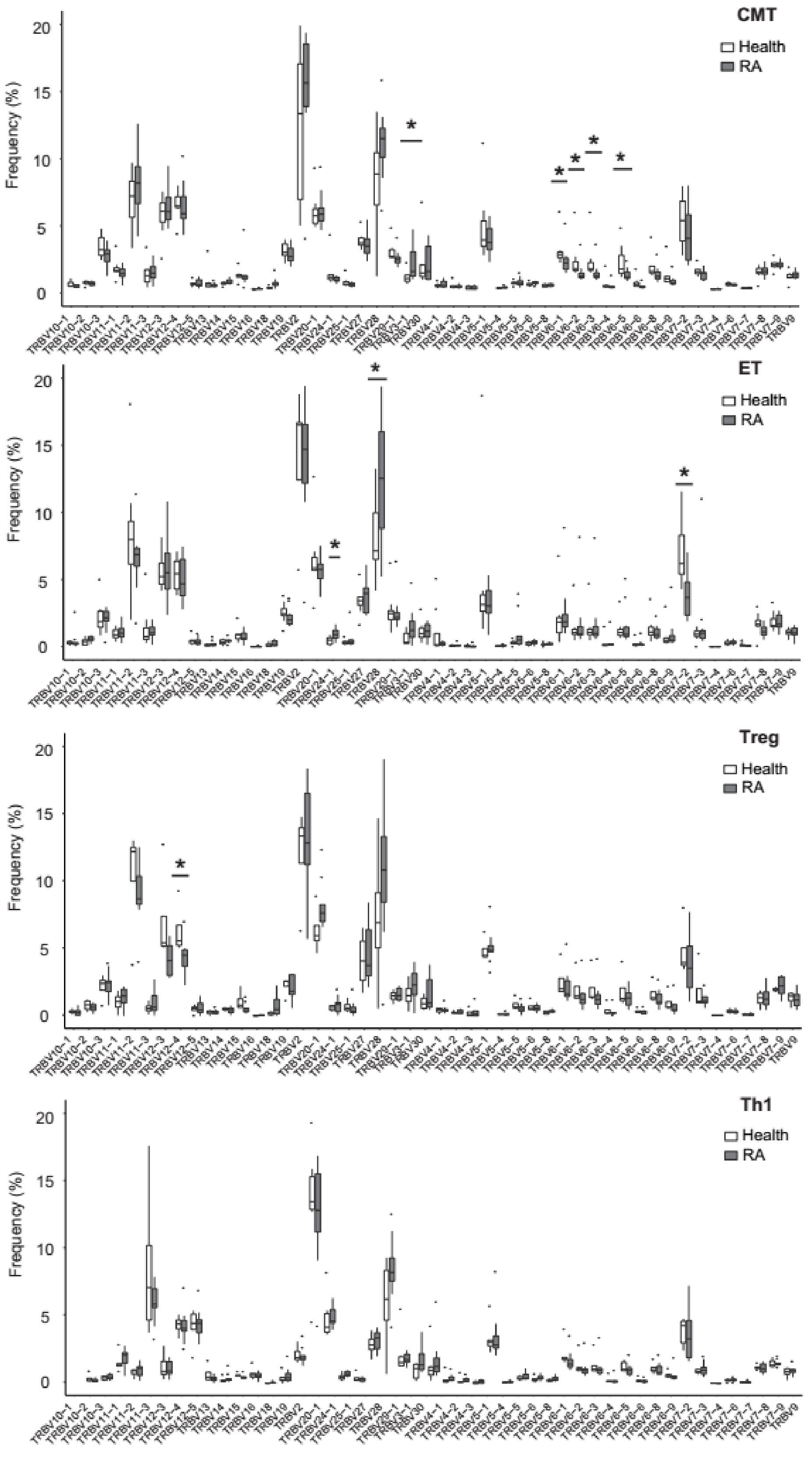


Figure S4. Different usages of V genes by CMT, ET, Treg and Th1 from RA and HC.

(Mann-Whitney U-test. *p<0.05.) (Data of CMT, ET, Treg, Th1 subsets sequenced successfully, as shown in tableS2, was analyzed.)

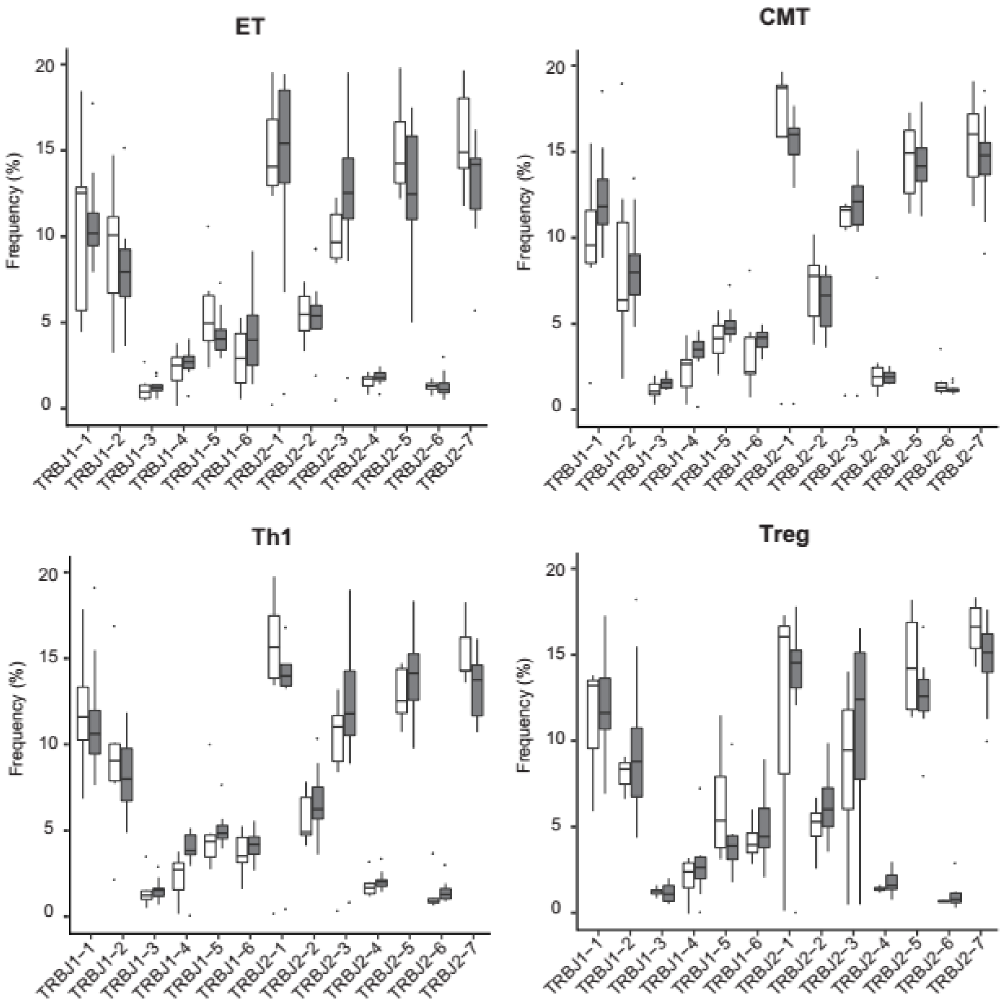


Figure S5. Different usages of J genes by ET, CMT, Th1 and Treg from RA and HC. (Data of ET, CMT, Th1, Treg subsets sequenced successfully, as shown in tableS2, was analyzed.)

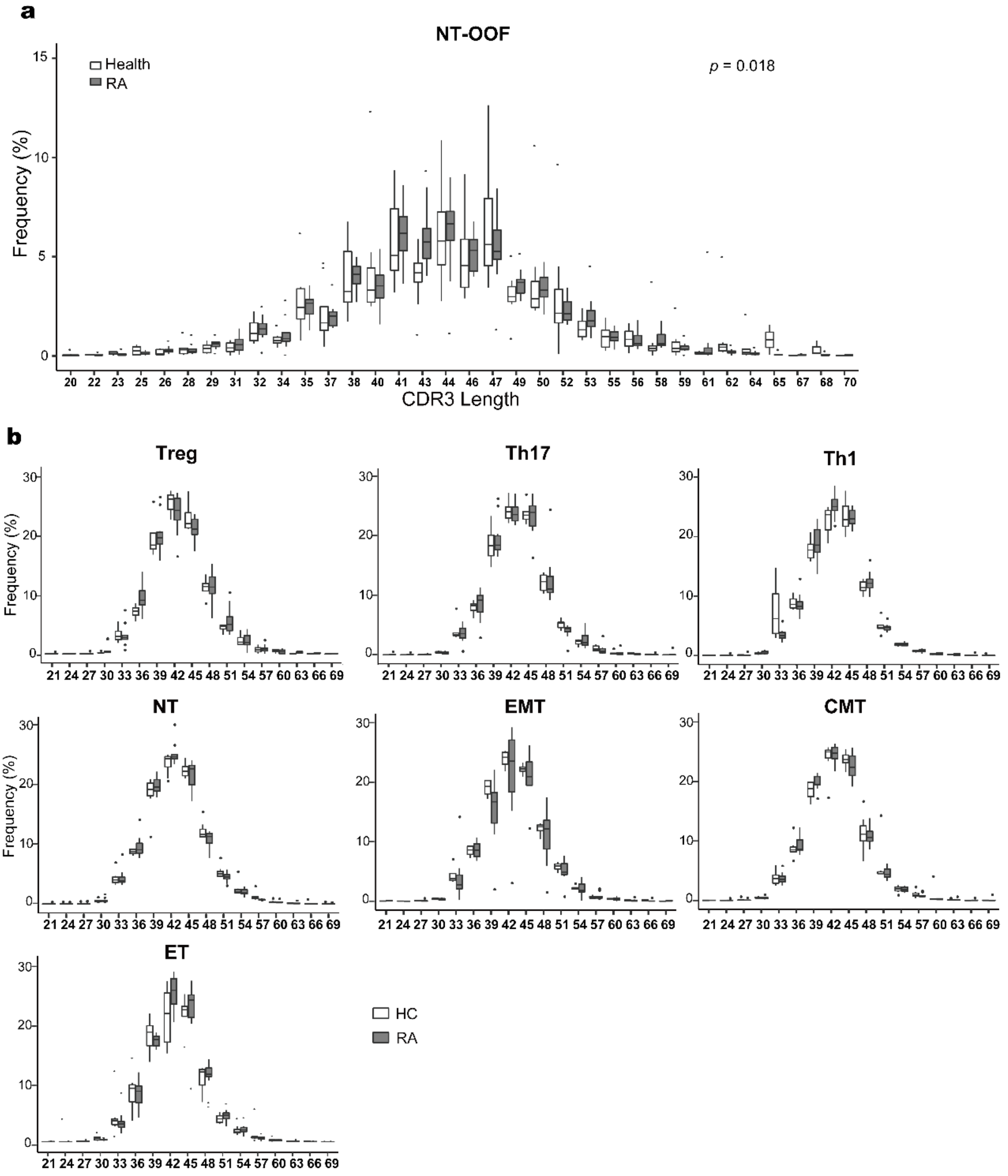


Figure S6. Pre-selection TCRB was reduced in RA patients than in HC.

(a) Distribution of CDR3 length of out-of-frame (OOF) sequences for TCR in RA and HC. (Kolmogorov–Smirnov test). (b) Distribution of CDR3 length of in-frame sequences for TCRs in RA and HC. (Data of all samples sequenced successfully, as shown in tableS2, was analyzed.)

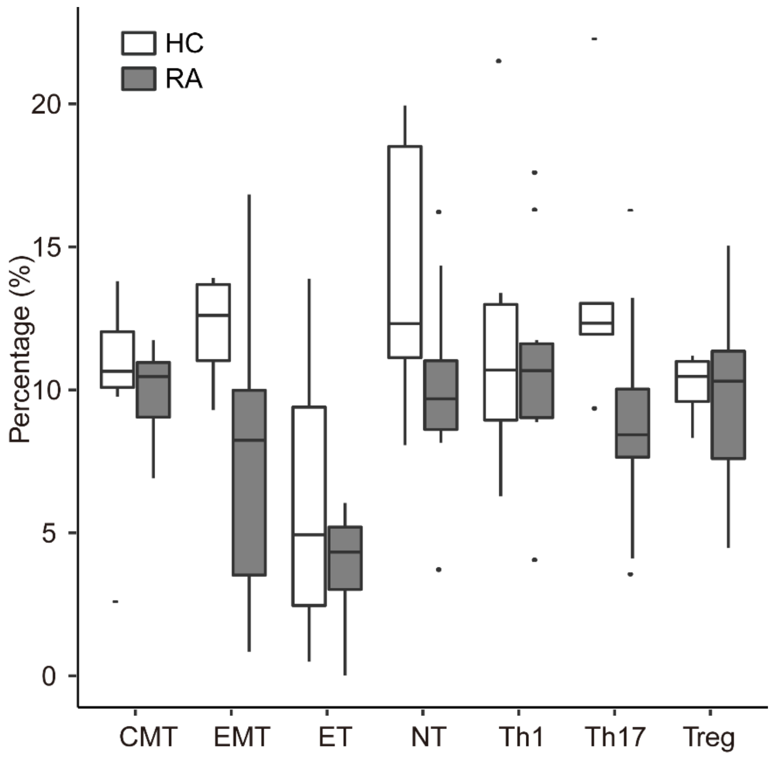

Figure S7. The D50 of each RA subset is similar to that of each HC.

The D50 index were calculated, no significant difference was observed in D50 indices.

(Data of all samples sequenced successfully, as shown in tableS2, was analyzed.)

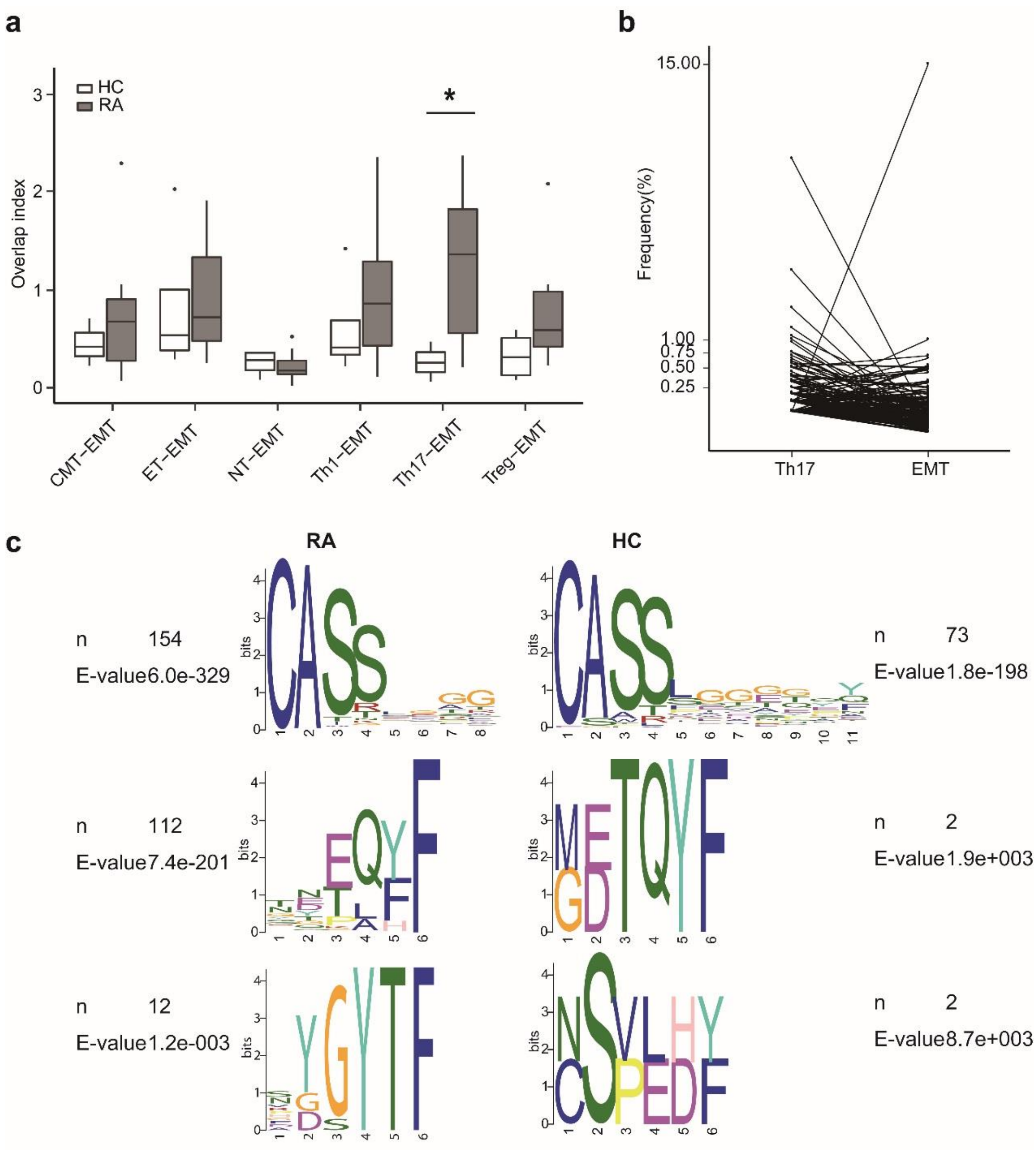


Figure S8. The overlapping index between EMT and other subsets.

(a) Overlapping index between EMT and other subsets were calculated, and CDR3s of EMT and Th17 from RA had more similarity in contrast to those from HC. (Mann-Whitney U-test. *p<0.05.) (the overlap index was calculated between subsets separated from the same donor, as shown in the tableS2.) (b) Overlapping clonotypes between Th17 and EMT had higher frequencies in Th17 than in EMT (126/167 conotypes). (c) These clonotypes in HC and RA present similarity motifs. (provided by

MEME)

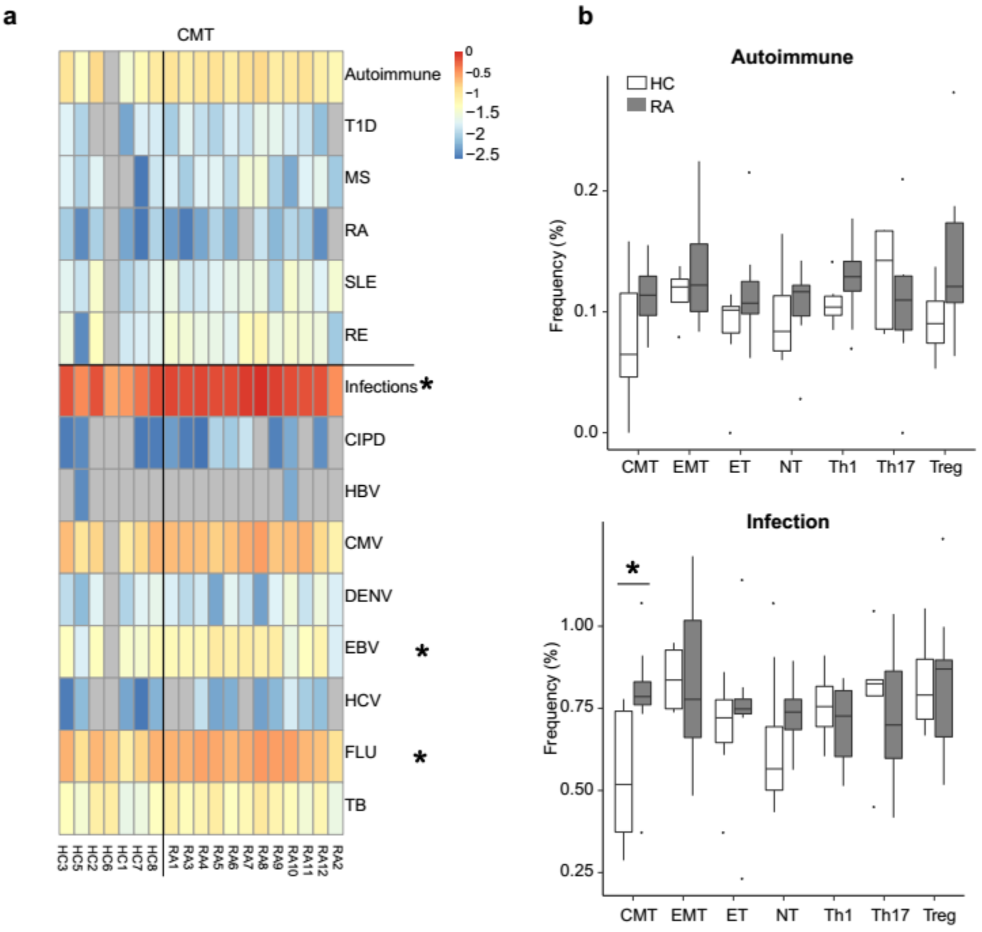


Figure S9 More clonotypes in CMT from RA are annotated as infection-related.

(a) Heatmap of disease-related clonotypes of CMT from RA and HC. Partly of clonotypes per specimen were identified in knowledge databases, and the frequency of them with same annotation in one donor were aggregated; the upper part in the plot is for autoimmune diseases, and the bottom is for infectious diseases; Log2(sum frequency) were plotted with different colors, and these sum frequencies annotated by the same disease were compared between RA and HC; (b) More clonotypes from CMT in RA were identified as infection-, EBV- and Flu-related sequences, however, no significant biases were observed in autoimmune disease-related sequences between RA and HC. (Mann-Whitney U-test. *p<0.05.) (Autoimmune: all autoimmune diseases recorded in the database; T1D: type 1 diabetes; MS: multiple sclerosis; RA: rheumatoid arthritis; SLE: systemic lupus erythematosus; RE: rasmussen encephalitis; infection: all infectious diseases recorded in the database; CIPD: Chronic inflammatory periodontal disease; HBV: hepatitis B virus; CMV: cytomegalovirus; DENV: dengue virus; EBV: Epstein–Barr virus; HCV: hepatitis C virus; FLU: influenza; TB: tuberculosis)(b, data of all samples sequenced successfully, as shown in tableS2, was analyzed.)
